# Defining strain variation in the maintenance of *Borrelia turicatae* during hyperparasitism and transovarial transmission in the tick vector

**DOI:** 10.64898/2026.07.30.741854

**Authors:** Serhii Filatov, Michael W. Curtis, Alexander R. Kneubehl, Winnie W. Kamau, Victor F. Figueroa, Brandon A. Hogland, Jon S. Blevins, Job E. Lopez

## Abstract

Tick-borne relapsing fever (TBRF) spirochetes are maintained in nature through vector-associated transmission routes primarily in argasid ticks. However, the extent to which strain-level differences influence the maintenance of TBRF spirochetes remains unclear. We evaluated two *Borrelia turicatae* strains, *Bt*–SSK1 and *Bt*–FCB, in *Ornithodoros turicata*. Both strains were acquired by female ticks, and after mating we unexpectedly observed that they were hyperparasitized by uninfected and unfed male *O. turicata. Bt*– SSK1 was maintained in males while *Bt–*FCB was not. Transovarial transmission (TOT) of *Bt*–SSK1 and *Bt*–FCB was also evaluated and striking differences were observed between strains. *Bt*–SSK1 was vertically maintained in F1 progeny but we failed to detect TOT of *Bt*–FCB. TOT of *Bt*–SSK1 occurred inefficiently in early ovipositions but increasing in later or delayed reproductive events, indicating a timing-dependent barrier. Using a *gfp*–expressing *Bt*–FCB strain, we show that failure of vertical transmission is associated with a lack of persistent oocyte colonization despite dissemination to ovarian tissues. These findings demonstrate strain-dependent differences in oocyte colonization and represent a critical bottleneck governing vertical transmission and persistence of relapsing fever spirochetes in tick populations.

**Importance:** Vector-borne pathogens rely on diverse transmission strategies to persist in nature, yet the biological factors that govern these processes remain poorly understood. In this study, we demonstrated that strains of *Borrelia turicatae* differ markedly in their ability to persist within *Ornithodoros turicata* ticks. We also discovered that male ticks frequently hyperparasitized engorged females, creating an unrecognized transmission route in which previously uninfected ticks can acquire, maintain, and transmitted the *B. turicatae*. We further showed phenotypic differences in vertical transmission and that it is temporally regulated and linked to successful colonization of developing oocytes, identifying a critical bottleneck in pathogen maintenance. Together, these findings revealed that strain-specific traits influence both horizontal maintenance within tick populations and vertical transmission to progeny. This work provides insight into the ecological and evolutionary processes that enable relapsing fever spirochetes to persist in nature.

## Introduction

The *Borreliaceae* family encompasses pathogenic spirochetes, which are the leading cause of bacterial vector-borne zoonoses worldwide (1, 2). Based on their clinical presentation, ecological characteristics, and evolutionary relationships, vector-borne spirochetes are further classified as Lyme disease-(LD) and relapsing fever-(RF) causing spirochetes (1). While most research has focused on LD-causing spirochetes, RF spirochetes are neglected yet distributed on five of seven continents and a major cause of illness and death if left untreated (2).

RF spirochetes are vectored by hard– (Ixodidae) and soft–bodied (Argasidae) ticks and the human body louse (*Pediculus humanus corporis*) (3). Argasid ticks in the genus *Ornithodoros* transmit nearly all species of RF spirochetes (4). In North America, there are two primary vectors of tick-borne relapsing fever (TBRF) spirochetes that impact human health. *Ornithodoros hermsi* vectors *Borrelia hermsii* and *Borrelia nietonii* (5), while *Borrelia turicatae* is maintained in *Ornithodoros turicata* ticks (6, 7). The specificity between *Ornithodoros* ticks and their associated *Borrelia* species shapes the geographic distribution and epidemiology of these pathogens.

The complex life cycles of *Ornithodoros* species highlight the importance of understanding how TBRF spirochetes exploit argasid tick biology. *Ornithodoros* species possess multiple nymphal instar stages, each of which requires a blood meal (8). The ticks can take years to mature into adults at which point they can blood feed numerous times (8). Laboratory studies have reported hyperparasitism in *Ornithodoros* species (9, 10), in which unfed ticks obtain a blood meal by feeding on recently engorged conspecifics (9, 10). Moreover, female *Ornithodoros* undergo multiple gonotrophic cycles (blood feeding and oviposition). While RF spirochetes have evolved to be vectored by argasid ticks, the intricacies of vector competence remain poorly understood.

A trait of TBRF spirochetes is transovarial transmission (TOT). Of the known TBRF *Borrelia*, TOT has been confirmed in 13 species (11, 12). In 1943, Gordon Davis’s seminal work with *B. turicatae* and *O. turicata* showed that this tick species maintained the spirochete long-term through TOT (13). He recovered *B. turicatae-*infected nymphs from a prairie dog burrow and a cottontail rabbit in Clark County, Kansas (KS). The nymphs were reared on a white rat, and one molted into an infected female tick. Over six years, he reared five successive generations of *O. turicata* from this female and determined the frequencies of *B. turicatae*-infected progeny across the cohorts. At the time, culture medium was unavailable to isolate and characterize *B. turicatae* strains, so the intricacies of TOT across the female tick’s life cycle are poorly understood.

Previously, we noted nuances in the maintenance of the *B. turicatae* strains (14, 15). The *B. turicatae* FCB (*Bt–*FCB) strain could be acquired by nymphal ticks during blood feeding, was transstadially maintained, and completed the tick–mouse transmission cycle (15). However, there was anecdotal evidence suggesting that the strain failed to undergo TOT. We also identified the *B. turicatae* SSK1 (*Bt-*SSK1) strain, which underwent TOT (14). Building on these preliminary observations, in this current study we investigated the maintenance of *Bt–*SSK1 and *Bt–*FCB in adult ticks and their offspring. We assessed *Bt–*SSK1 and *Bt–*FCB acquisition by female *O. turicata,* noted hyperparasitism by male ticks during breeding, and characterized TOT of these two strains over successive gonotrophic cycles. We observed nuances in *O. turicata* reproduction and phenotypic differences in TOT between *Bt–*SSK1 and *Bt–*FCB. Colonization of female *O. turicata* was further investigated by generating *Bt–*FCB producing the green fluorescent protein (*Bt–*FCB*–*GFP). Together, our findings provide new insights into the maintenance of these understudied zoonotic spirochetes.

## Results

### Evaluation of *O. turicata-*KS and TX colonies

We used two colonies of *O. turicata* that originated from Kansas (KS) and Texas (TX). It is not possible to distinguish between male and female *O. turicata* until nymphs molt into adults and the genital aperture appears. Consequently, when we selected *O. turicata–*KS females from the colony their reproductive status was unknown because they were mixed with males and they likely had already mated.

For the *O. turicata–*TX colony, these ticks originated from a female collected in a park in Austin, Texas (14). The F1 progeny from this female were determined to be free of *B. turicatae* by feeding ticks at the nymphal stage through adulthood on mice, and spirochetes were never visualized in the blood by dark field microscopy. Thirty days after tick feedings, immunoblotting was performed and mice failed to seroconvert to *B. turicatae* protein lysates. These findings further supported an absence of infection in mice and that the tick colonies were free of *B. turicatae*.

We also controlled for the reproductive status of *O. turicata–*TX. We monitored late instar stage nymphs daily and observed that males emerged first, at which point they were removed from the cohort. As females emerged from the final nymphal stage, they were removed and housed separately to ensure that they were unmated (virgin) ticks. These *O. turicata-*TX females served as the source of adults used in the studies below.

### Acquisition of *Bt-*SSK1 and *Bt-*FCB by female *O. turicata*

Figure 1 shows the experimental approach used to infect *O. turicata–*KS and *O. turicata–*TX and evaluate the tick–mammalian transmission cycle and TOT of *Bt–*SSK1 and *Bt–*FCB. Mice that were infected by intraperitoneal inoculation were spirochetemic the following day, while animals infected by tick bite became spirochetemic within five days. At the spirochete acquisition blood meal (Fig. 1), spirochete densities were between 2.1 x 10^4^ to 1.5 x 10^5^ bacteria per ml of murine blood, and all *O. turicata–*KS and *–*TX female ticks fed to repletion. After blood feeding, *O. turicata–*KS and *–*TX females successfully mated with an individual male that originated from the same colony (Fig. 1).

**Figure 1.**
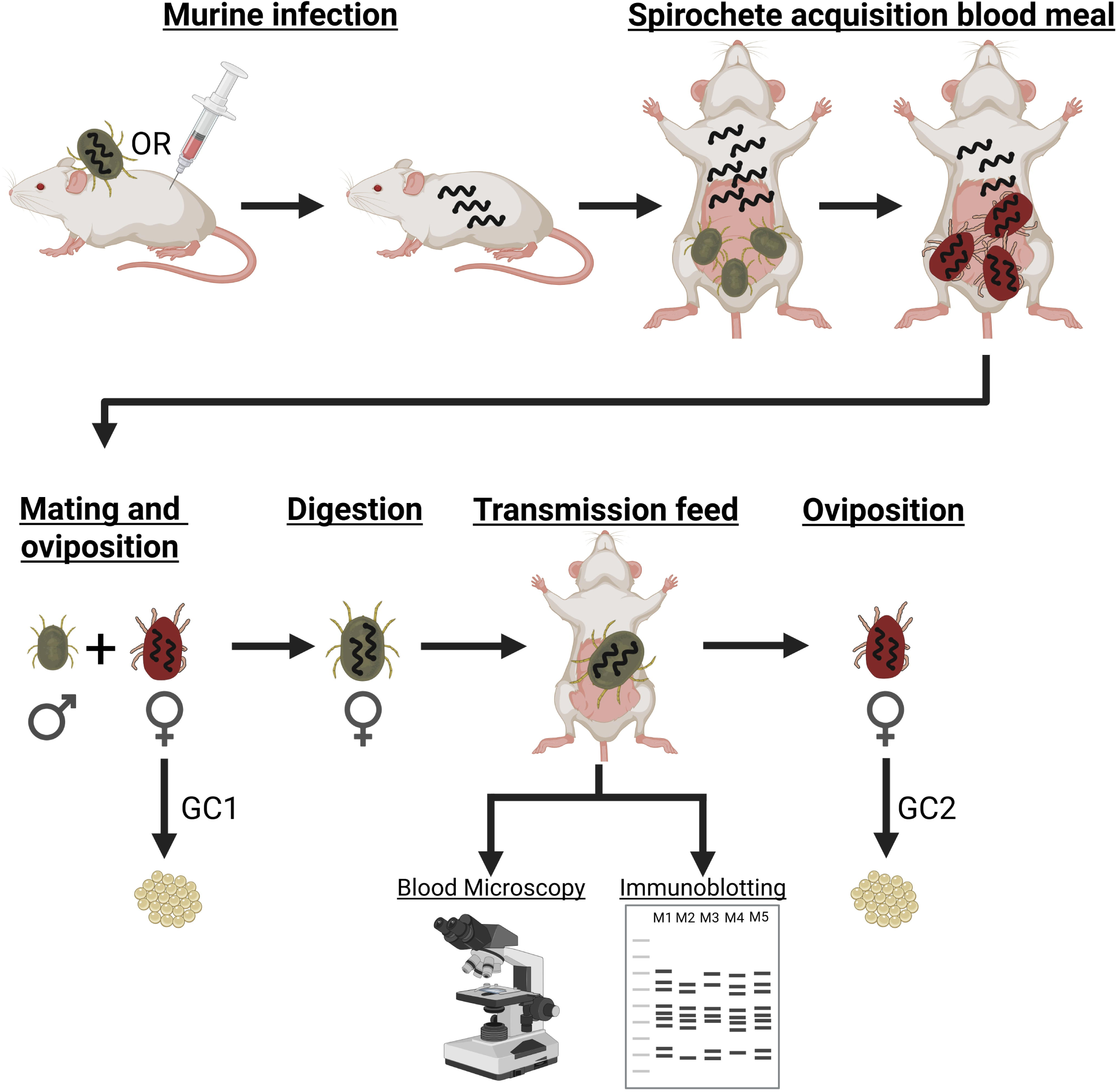
Experimental approach to infect *O. turicata* with *B. turicatae* and to assess tick colonization and reproduction. Mice were infected by needle inoculation or tick bite with one of the two *B. turicatae* strains (SSK1 or FCB). The following day mice were confirmed to be infected by dark field microscopy and qPCR (not shown), and a spirochete acquisition blood meal was performed. Engorged females were allowed to mate with male ticks and females laid eggs in their first gonotrophic cycle (GC1). After digesting the blood meal, female infection was validated by allowing a single tick to feed on a mouse and infection was determined by dark field microscopy to visualize live spirochetes. Infection was also assessed by immunoblotting to detect seroconversion to *B. turicatae* proteins lysates 30 days after the transmission feed. Aslo shown is the second gonotrophic cycle (GC2). This image was generated with BioRender.

### Hyperparasitism of female ticks extended the life cycle of *Bt–*SSK1

After the acquisition blood meals, we observed feeding scars on females or engorged male ticks, which revealed that female *O. turicata* had been hyperparasitized by the male tick they were partnered with for reproduction. We first observed discernable scars on *O. turicata–*KS females (Fig. 2). Closer observation indicated that nine out of 10 *O. turicata–*KS female ticks that were colonized by *Bt–*SSK1 had feeding scars, while three of five females colonized with *Bt–*FCB had scars (Table 1). For *O. turicata*–TX ticks, hyperparasitism was closely monitored after pairing females with males. Three out of five males engorged on female ticks within the first 24 hours of pairing, irrespective of whether they were colonized with *Bt*–SSK1 or *Bt*–FCB (Table 1).

**Figure 2.**
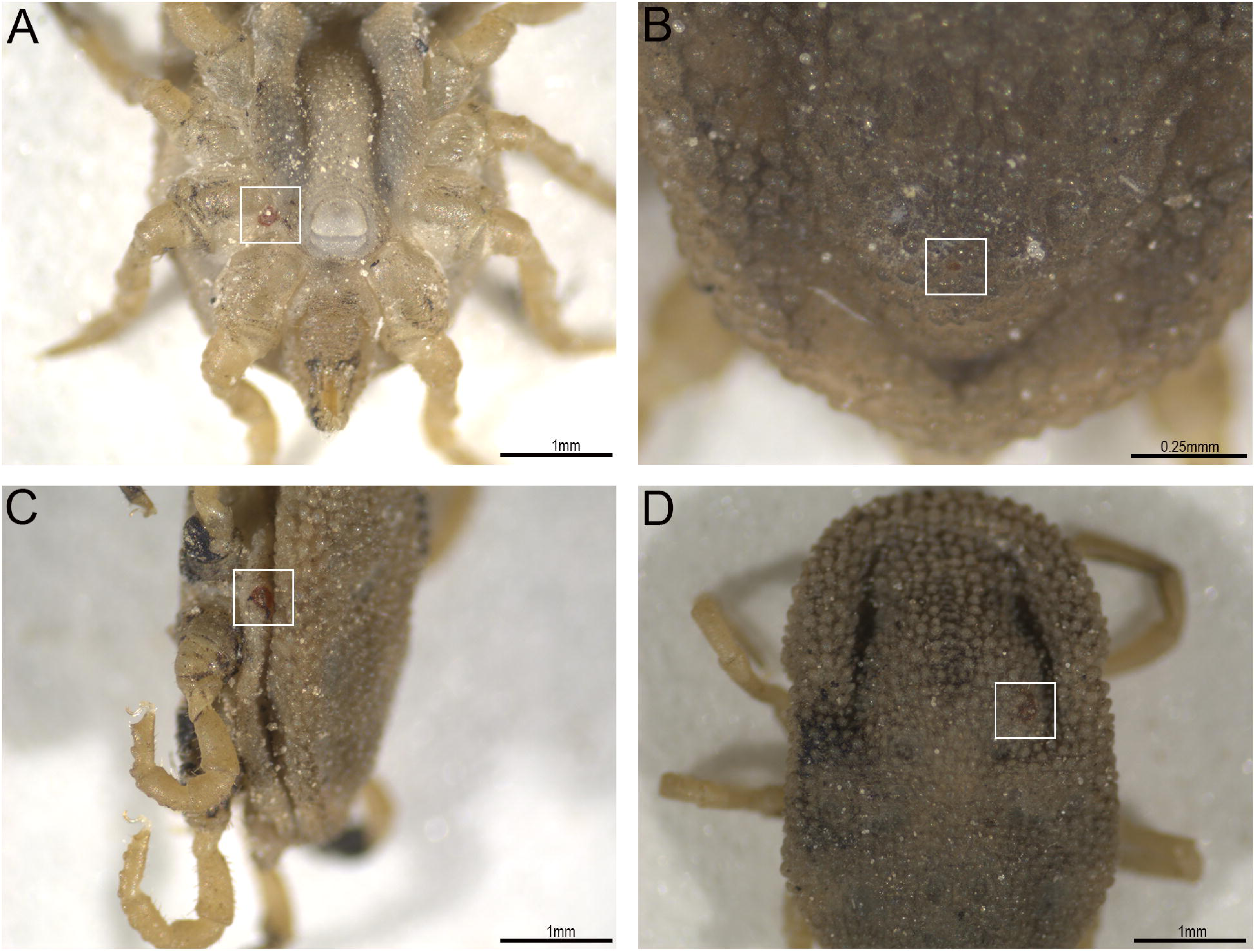
Examples of hyperparasitism in a group of *O. turicata* females infected with the SSK1 strain. In most cases, the scars left by the male’s mouthparts (white square) remained easily identifiable months after attachment. Shown in (A) is a female on the ventral side, between left coxae I and II, while in (B) there is a female with a scar in the anterior-central region of the dorsum. Shown in (C) is a female that was parasitized in the lateral region, between the right coxae II and III, while (D) shows a female bearing a scar in the central region of the dorsum. A scale bar is shown in the lower right of each panel.

**Table 1.** Hyperparasitism of female *O. turicata* by male ticks.

| *B. turicatae* strain | *O. turicata*-KS |  |  | *O. turicata*-TX |  |
| --- | --- | --- | --- | --- | --- |
|  | SSK1 | SSK1 | FCB | SSK1 | FCB |
| Mouse inoculation route | IP | tick bite | IP | IP | IP |
| Female scar | 4/5 | 5/5 | 3/5 | 3/5 | 3/5 |
| Male transmission | 4/5 | 3/4 <sup>A</sup> | 0/5 | 2/5 | 0/5 |

A: One male tick died before blood feeding on a mouse

Since hyperparasitism was considered uncommon for *O. turicata* (16), we investigated whether *Bt–*SSK1 and *Bt–*FCB could infect mice by tick transmission after male ticks digested their blood meals (Table 1). Infected animals were spirochetemic within five days after blood feeding male ticks, as determined by dark field microscopy. *O. turicata–*KS males delivered an infectious dose of *Bt–*SSK1 to seven of nine mice (Table 1). Moreover, *O. turicata–*TX males infected two of five mice with *Bt–*SSK1. We failed to detect *Bt–*FCB infection in murine blood after feeding male *O. turicata–*KS or *–* TX ticks twice on mice, and none of these animals seroconverted to *B. turicatae* protein lysates (Fig. 3). These findings indicated that *Bt–*FCB failed to establish an infection in mice after being fed on by male ticks that originally hyperparasitized on engorged female ticks. Unfortunately, male ticks died before we could perform qPCR to evaluate whether they were colonized with *Bt–*FCB.

**Figure 3.**
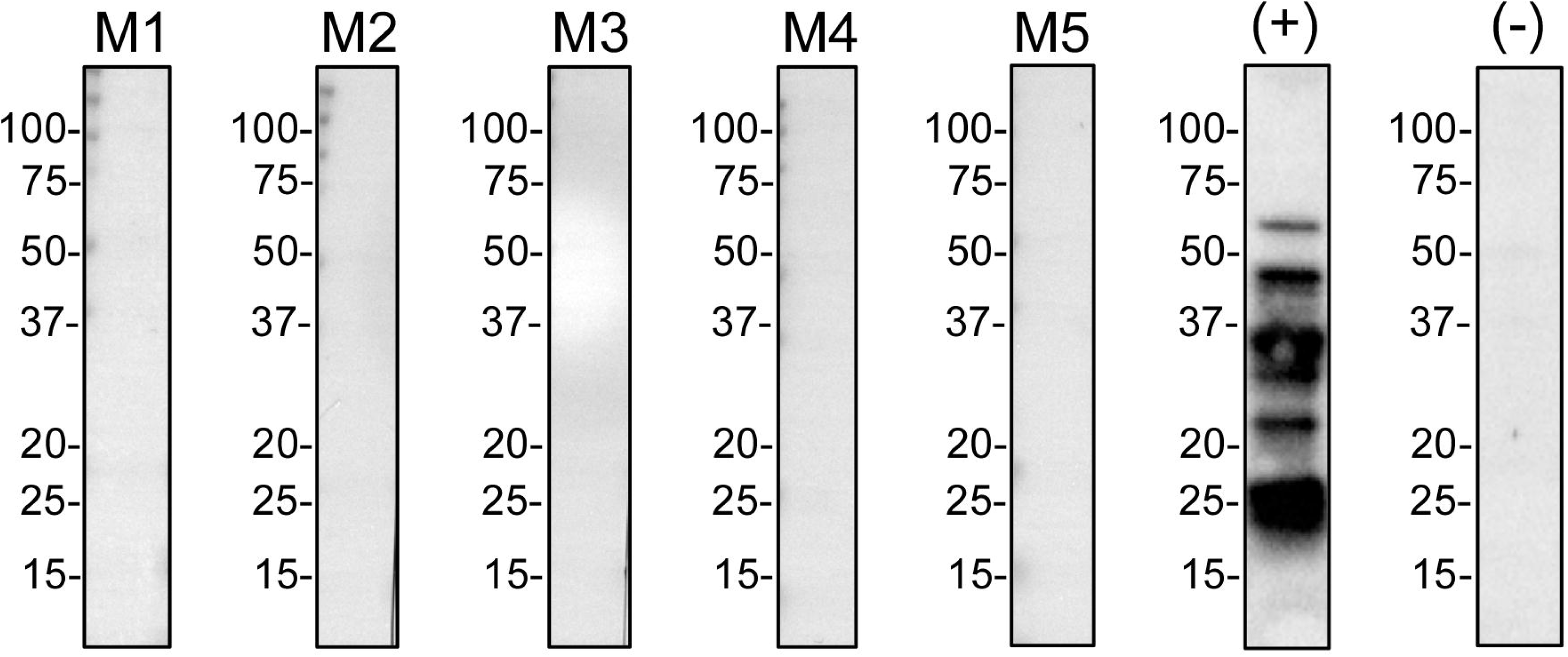
Serological responses to *B. turicatae* protein lysates using serum samples from mice that were exposed to male *O. turicata* ticks*. B. turicatae* proteins lysates were used for immunoblots. Shown are immunoblots using serum samples from five mice exposed to male ticks and represent the remainder of the animals that were seronegative. A positive (+) and negative (-) control are also shown. Molecular masses are shown in kDa to the left of each immunoblot.

By noting feeding scars on females, recording the number males that engorged on female *O. turicata*, and evaluating male transmission to mice, we determined that 19 of 24 (78.2%) female ticks were hyperparasitized. We determined that hyperparasitism did not impact female viability because within 30 days they had digested the blood meal and remained alive. After observing hyperparasitism, we mated a cohort of virgin *O. turicata-*TX female ticks prior to feeding them on a *Bt–*SSK*–*infected mouse, and these served as a control for TOT experiments.

### Female ticks colonized with *Bt–*SSK1 or *Bt–*FCB delivered an infectious dose of spirochetes to mice after blood feeding

Once females digested the *B. turicatae* acquisition blood meal, we determined whether they could deliver an infectious dose of *Bt-*SSK1 and *Bt–*FCB to naïve mice (transmission feed in Fig. 1). Regardless of the tick colony used, both *Bt–*SSK1 and *Bt–* FCB were infectious by tick bite (Table 2). Furthermore, hyperparasitism did not impact the ability of female ticks to infect mice during the transmission blood meal compared to the non-hyperparasitized controls (Table 2).

**Table 2.** Assessment of the tick-mammalian transmission cycle of *Bt-*SSK1 and *Bt-*FCB in female *O. turicata*.

| *B. turicatae* strain | *O. turicata*-KS |  |  | *O. turicata*-TX |  |  |
| --- | --- | --- | --- | --- | --- | --- |
|  | SSK1 | SSK1 | FCB | SSK1 | SSK1 | FCB |
| Mouse inoculation | IP | Tick bite | IP | IP | Tick bite | IP |
| Hyperparasitism | Yes | Yes | Yes | Yes | No<sup>A</sup> | Yes |
| Murine infection after feeding female ticks | 5/5 | 4/4 | 4/4 | 5/5 | 5/5 | 3/5 |

A: These female ticks were mated prior to blood feeding

### The offspring of *Bt–*SSK1- and *Bt–*FCB–colonized female ticks differed in their ability to infect mice after blood feeding

In Figure 4 we show the experimental approach to assess TOT of *Bt–*SSK1 and *Bt–*FCB from the offspring of female *O. turicata–*KS and *–*TX ticks. Once eggs hatched, the larvae and first instar stage nymphs were successfully reared on mouse pups (Fig. 4). At the second instar stage, we pooled 15 offspring ticks from a given female, fed them on individual mice, and observed marked differences in murine infection between the *Bt–*SSK1 and *Bt–*FCB (Table 3).

**Figure 4.**
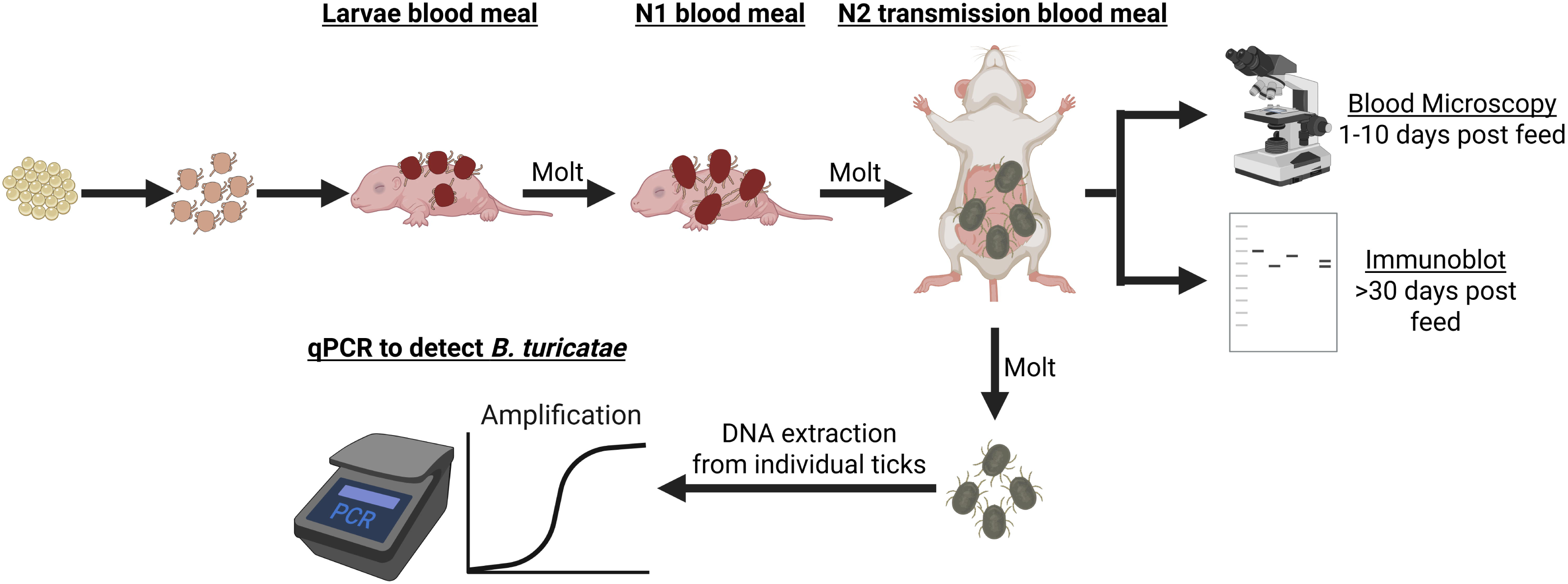
Experimental approach to assess TOT and filial infection rates of *B. turicatae* in the vector. Progeny from each gonotrophic cycle were reared as larvae and fist instar stage nymphs (N1) on mouse pups. At the second instar stage (N2) ticks were pooled for test-feeding on mice to assess vertically acquired *B. turicatae* infection. Following their molt, genomic DNA was extracted from individual ticks for qPCR to determine filial infection rates. This image was generated with BioRender.

**Table 3.** Evaluating murine infectivity of *Bt-*SSK1 and *Bt-*FCB after feeding the offspring of infected female ticks.

**No. of mice infected after feeding 2nd instar stage nymphs<sup>A</sup> from:**
| Colony | *Bt* strain | *Bt* acquisition | GC | Female 1 | Female 2 | Female 3 | Female 4 | Female 5 | Total mice infected |
| --- | --- | --- | --- | --- | --- | --- | --- | --- | --- |
| *Ot*-KS | SSK1 | IP infected mice |  |  |  |  |  |  |  |
|  |  |  | 1 | + (4/4) | - (0/4) | + (3/3) | - (0/3) | + (1/1) | 8/15 (53%) |
|  |  |  | 2 | + (4/4) | + (4/4) | + (4/4) | + (4/4) | NA | 16/16 (100%) |
| *Ot*-KS | SSK1 | Tick bite<br>infected mice | 1 | NA | - (0/4) | + (4/4) | + (4/4) | + (4/4) | 12/16 (75%) |
|  |  |  | 2 | NA | + (1/1) | + (4/4) | + (4/4) | + (4/4) | 13/13 (100%) |
| *Ot*-KS | FCB | IP infected mice | 1 | - (0/4) | - (0/4) | - (0/4) | - (0/4) | NA | 0/16 (0%) |
|  |  |  | 2 | - (0/4) | - (0/4) | - (0/4) | - (0/4) | NA | 0/16 (0%) |
| *Ot*-TX | SSK1 | IP infected mice | 1 | - (0/4) | - (0/4) | - (0/4) | - (0/4) | - (0/4) | 0/20 (0%) |
|  |  |  | 2 | + (4/4) | + (4/4) | NA | - (0/4) | + (4/4) | 12/16 (75%) |
| *Ot*-TX<sup>B</sup> | SSK1 | Tick bite<br>infected mice | 1 | - (0/4) | - (0/4) | - (0/4) | - (0/4) | - (0/4) | 0/16 (0%) |
|  |  |  | 2 | + (4/4) | + (2/4) | + (4/4) | + (4/4) | + (4/4) | 18/20 (90%) |
| *Ot*-TX | FCB | IP infected mice | 1 | - (0/4) | - (0/4) | - (0/4) | - (0/4) | - (0/4) | 0/20 (0%) |
|  |  |  | 2 | - (0/4) | - (0/4) | - (0/4) | - (0/4) | - (0/4) | 0/20 (0%) |

**A:** 15 offspring ticks fed upon per mouse;
**B:** Offspring from non-hyperparasitized female ticks
**GC:** Gonotrophic cycle
**NA:** Female died

Offspring arising from the first gonotrophic cycle of *O. turicata–*KS females colonized with *Bt–*SSK1 successfully infected 53% to 75% of mice with spirochetes (Table 3). Offspring from the second gonotrophic cycle of the same females successfully infected 100% of mice with *Bt–*SSK1 (Table 3). We failed to detect murine infection after feeding the offspring of adult female *O. turicata–*KS ticks colonized with *Bt–*FCB (Table 3).

After feeding the offspring of *O. turicata-*TX female ticks, we again observed differences in murine infection between *Bt–*SSK1 and *Bt–*FCB. Offspring from the first gonotrophic cycle failed to infect mice with *Bt–*SSK1. However, when we fed offspring from the second gonotrophic cycle, 75% to 90% of animals became infected (Table 3). We found no differences in the ability of offspring from hyperparasitized and non-hyperparasitized *O. turicata–*TX females colonized with *Bt–*SSK1 to successfully infect mice (Table 3). We also failed to detect infection in mice after feeding the offspring of *Bt–*FCB colonized females.

Evaluation of mouse antibody responses to *B. turicatae* protein lysates demonstrated that animals in which *Bt*–SSK1 and *Bt*–FCB were undetectable by dark-field microscopy also failed to seroconvert. This indicated that an infection beneath the threshold detectable by dark field microscopy did not occur. Our work established that *Bt–*SSK1 was infectious to mice from female *O. turicata*–KS and –TX ticks and their offspring, while *Bt–*FCB was only infectious to animals from female ticks. Furthermore, there was no difference in transmission from the offspring of hyperparasitized and non-hyperparasitized females. This suggested that rupturing of the midgut did not facilitate spirochete dissemination to developing oocytes and enhance TOT.

### Molecular detection of *Bt-*SSK1 and *Bt-*FCB in the offspring of *O. turicata-*KS ticks

Twenty offspring nymphs from each gonotrophic cycle were randomly selected from each *O. turicata*–KS female, and qPCR identified phenotypic differences in filial infection rates between *Bt–*SSK1 and *Bt–*FCB (Table 4). Regardless of how female ticks acquired *B. turicatae,* filial infection rates were significantly higher in the second gonotrophic cycle with a large effect size (Cliff’s δ = -0.47, 95% CI: -0.65 to -0.4, averaged across replications). In contrast, all 160 progeny from FCB-infected females tested negative for *Bt-*FCB by qPCR (Table 4). These results indicated that *Bt–*SSK1 is vertically maintained by *O. turicata* while *Bt–*FCB fails to undergo TOT.

**Table 4.** Filial infection frequencies from O. turicata-KS female ticks.

| Strain | *Bt* acquisition | GC | Filial infection rates from (% , 95% CI): |  |  |  |  |
| --- | --- | --- | --- | --- | --- | --- | --- |
|  |  |  | Female 1 | Female 2 | Female 3 | Female 4 | Female 5 |
| SSK1 | Intraperitoneal |  |  |  |  |  |  |
|  |  | GC1 | 6/20<br>(30%, 13-54%) | 0/20<br>(0%, 0-20%) | 7/20<br>(35%, 16-59%) | 0/20<br>(0%, 0-20%) | 3/18<br>(16.7%, 4-42%) |
|  |  | GC2 | 16/20<br>(80%, 59-93%) | 14/20<br>(70%, 46-87%) | 14/20<br>(70%, 46-87%) | 12/20<br>(60%, 36-80%) | NA |
|  |  | p-value* | 0.004 | 3.34E-06 | 0.056 (NS) | 4.51E-05 |  |
| SSK1 | Tick bite |  |  |  |  |  |  |
|  |  | GC1 | NA | 0/20<br>(0%, 0-20%) | 1/20<br>(5%, 0.26-27%) | 2/40<br>(5%, 0-20%) | 1/20<br>(5%, 0.26-27%) |
|  |  | GC2 | NA | 5/20<br>(25%, 9-41%) | 16/20<br>(80%, 56-93%) | 4/40<br>(10%, 0-20%) | 17/20<br>(85%, 61-96%) |
|  |  | p-value* |  | 0.047 | 2.21E-06 | 0.6752 (NS) | 4.06E-07 |
| FCB | Intraperitoneal |  |  |  |  |  |  |
|  |  | GC1 | 0/20<br>(0%, 0-20%) | 0/20<br>(0%, 0-20%) | 0/20<br>(0%, 0-20%) | 0/20<br>(0%, 0-20%) | NA |
|  |  | GC2 | 0/20<br>(0%, 0-20%) | 0/20<br>(0%, 0-20%) | 0/20<br>(0%, 0-20%) | 0/20<br>(0%, 0-20%) | NA |
|  |  | p-value* | NC | NC | NC | NC | NC |
|  | .. |  |  |  |  |  |  |

**GC:** Gonotrophic cycle
**NA:** Not applicable, tick died
**NC:** Not calculated
\* p-value between GC1 and GC2

### TOT of *Bt–*SSK1 is linked to the reproductive kinetics of its vector

We observed frequent gonotrophic dissociation in *O. turicata–*KS and *–*TX females, regardless of if they were colonized with *Bt–*SSK1 or *Bt–*FCB. In total, 18 of 28 females (64.3%) exhibited irregularities in oviposition either in first or second gonotrophic cycle, laying a second smaller batch of eggs 30 to 60 days later without requiring an additional blood meal. Since the second oviposition of the first gonotrophic cycle represents an intra-cycle deviation that has not been previously reported for *O. turicata* (16, 17), we excluded those ticks from the statistics analysis in Table 4.

We further evaluated gonotrophic dissociation and quantified filial infection rates from *O. turicata–*KS female ticks 2 and 4 (Table 4). These females failed to produce *Bt–* SSK1 infected progeny in the first oviposition of their first gonotrophic cycle but exhibited gonotrophic dissociation. Interestingly, offspring from the second oviposition of the first gonotrophic cycle were infected with spirochetes (Fig. 5 A and B). These females were blood fed again and vertically transmitted *Bt–*SSK1 in the second gonotrophic cycle (third oviposition) at filial infection rates similar to the preceding batch (Figure 5 A and B). Our findings indicate that the temporal lag between ovipositions within the same gonotrophic cycle was sufficient for vertical transmission to occur.

**Figure 5.**
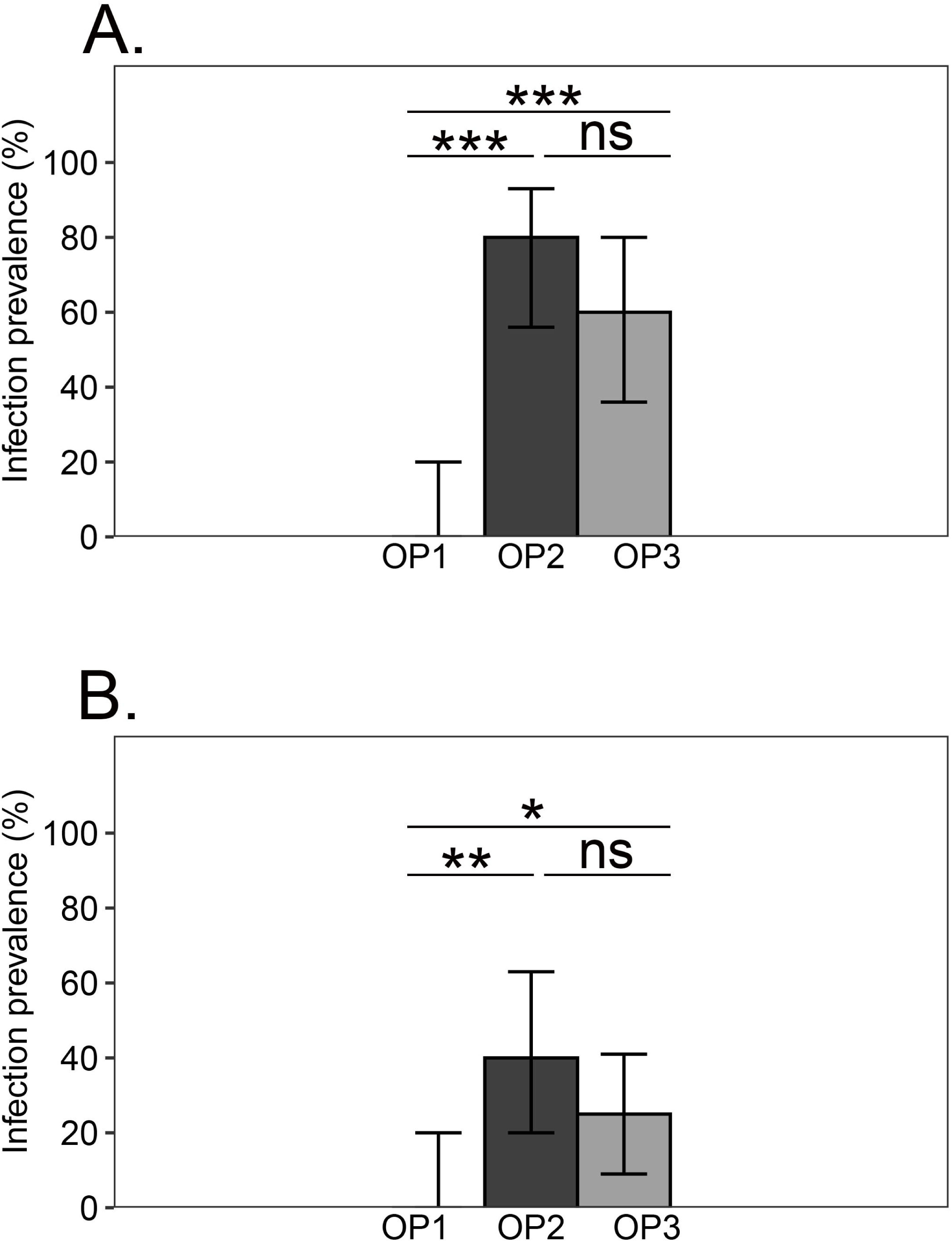
Filial infection rates of *Bt-*SSK1 in two *O. turicata* females exhibiting gonotrophic dissociation. Ovipositions 1 and 2 (OP 1 and OP2) occurred in the first gonotrophic cycle (A). The third oviposition (OP3) occurred in the subsequent gonotrophic cycle. Similar findings were observed in the offspring of another female tick infected with *Bt-*SSK1 (B). Fisher’s exact test was applied to compare FIRs between the ovipositions. Error bars represent the Wilson 95% confidence interval. ns = p > 0.05; * = p < 0.05; ** = p < 0.01; *** = p < 0.001.

### Generation of *Bt–*FCB–*gfp*

We further investigated female tick colonization by *Bt–*FCB through the generation of spirochetes that stably express *gfp*. We created a *trans* integration construct for *B. turicatae* similar to that developed for *B. duttonii* (18). A 495–bp intergenic region between two convergently transcribed genes in the 158-kb linear megaplasmid (e.g., *bta019* and *bta020*) was chosen as an integration site. Regions flanking the integration site were amplified, and a cassette containing *aph*[3’]*IIIa* and P*flaB*-*gfp* was ligated between them, creating the integration construct pBtKan-*gfp* (Fig. 6 A). This construct was subsequently transformed into *Bt–*FCB, resulting in *Bt–*FCB– *gfp*. Confirmation of the integration in *Bt–*FCB–*gfp* was accomplished by PCR to amplify across the insertion site between *bta019* and *bta020,* as well as internal regions of *gfp*, *aph*[3’]-*IIIa*, and *flaB* (Fig. 6 B). *Bt–*FCB–*gfp* screened positive for *gfp* and *aph*[3’]-*IIIa*, while *Bt–*FCB did not. The PCR amplification across the integration site yielded amplicons of the expected sizes: 314 bp and 2,341 bp for *Bt–*FCB and *Bt–*FCB–*gfp,* respectively. Additionally, PCR for *flaB*, serving as an amplification control, produced the correct-sized products for *Bt–*FCB and *Bt–*FCB–*gfp*. Fluorescence microscopy confirmed GFP production in *Bt–*FCB–*gfp* (Fig. 6 C). Overall, these results verify the integration of the *aph*[3’]*IIIa*-P*flaB*-*gfp* cassette into the chosen site within the 158-kb linear megaplasmid.

**Figure 6.**
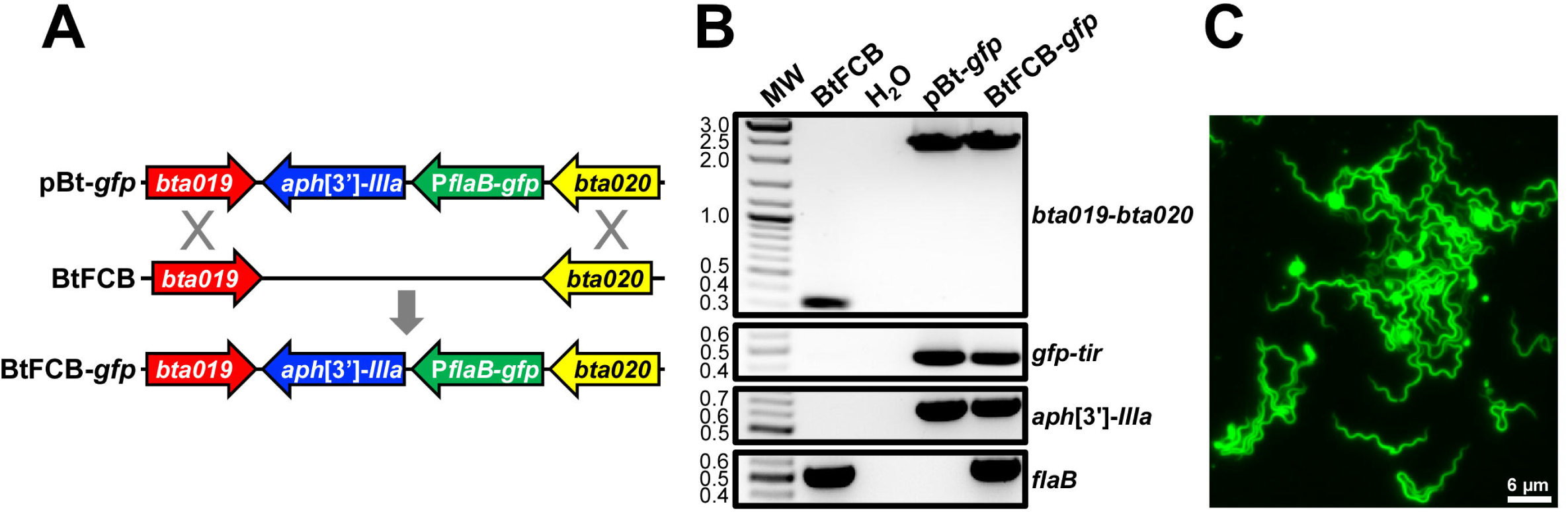
Integration of Kan-*gfp* into the 158-kb linear megaplasmid of FCB. In (A), we show the approach used to generate *Bt*-FCB-*gfp*. *Bt-*FCB was transformed with an allelic exchange construct (pBtKan-*gfp*) to insert P*flaB-gfp* and *aph*[3′]-*IIIa* cassettes into the 158-kb linear megaplasmid, generating *Bt*-FCB-*gfp*. In (B), PCRs were performed on gDNAs isolated from *Bt*-FCB and *Bt*-FCB-*gfp* transformants. Plasmid pBtKan-*gfp* was included as a positive amplification control, and PCRs were also performed with no template (H_2_O) as a purity control. Diagnostic PCRs, amplicons identified on the right, amplified a region flanking the *bta019-bta020* integration site (wild-type = 314 bp and integrant = 2,341 bp), as well as internal regions of *gfp* (456 bp), *aph[3’]-IIIa* (624 bp), or *flaB* (531 bp). “MW” denotes the DNA standard, and numbers to the left indicate molecular weight in kb. In (C), *gfp*-expressing *Bt*-FCB-GFP were visualized by fluorescence microscopy.

### Evaluation of *Bt–*FCB–*gfp* colonization in female *O. turicata*

Since *Bt–*FCB failed to infect progeny of colonized female ticks, we investigated whether the strain invades female reproductive tissues. To test this, mice were needle inoculated with 1 x 10^7^ *Bt–*FCB–*gfp* and became spirochetemic the following day with 3.98 x 10^5^ to 1 x 10^6^ bacteria per ml of murine blood (Table 5). Nine of 10 virgin female *O. turicata*–KS fully engorged on mice and were successfully mated. A partially engorged female was excluded from the analysis.

**Table 5.** Detection of Bt-FCB-gfp spirochetes in the midgut, salivary glands, and ovaries at time points post acquisition.

| Female | Spirochetes densities per ml of blood at acquisition | Female tick transmission | GC | Tick dissection following *Bt*-FCB-*gfp* acquisition (months) | Midgut | Salivary glands | Ovary |
| --- | --- | --- | --- | --- | --- | --- | --- |
| 1 | $3.98 \times 10^5$ per ml | N/A | 1 | 1 | + | + | - |
| 2 | $3.98 \times 10^5$ per ml | N/A | 1 | 1 | + | + | - |
| 3 | $3.98 \times 10^5$ per ml | + | 2 | 10 | + | + | + |
| 4 | $3.98 \times 10^5$ per ml | + | 2 | 10 | + | + | + |
| 5 | $3.98 \times 10^5$ per ml | + | 2 | 10 | + | + | + |
| 6 | $3.98 \times 10^5$ per ml | + | 2 | 10 | - | - | - |
| 7 | $1 \times 10^6$ per ml | - | 2 | 16 | + | + | + |
| 8 | $1 \times 10^6$ per ml | - | 2 | 16 | + | + | + |
| 9 | $1 \times 10^6$ per ml | - | 2 | 16 | - | - | - |
| Total |  | 4/7 |  |  | 7/9 | 7/9 | 5/9 |

**GC:** Gonotrophic cycle

The midguts, salivary glands, and reproductive tissues of female *O. turicata* were assessed for colonization by *Bt–*FCB–*gfp* at different time points post acquisition (Fig. 7). Two females were evaluated after the first gonotrophic cycle (Table 5). These ticks had started to oviposit when we dissected them, and fluorescing spirochetes were visualized in the midgut and salivary glands of both ticks, but we failed to detect *Bt–*FCB–*gfp* in reproductive tissues (Table 5).

**Figure 7.**
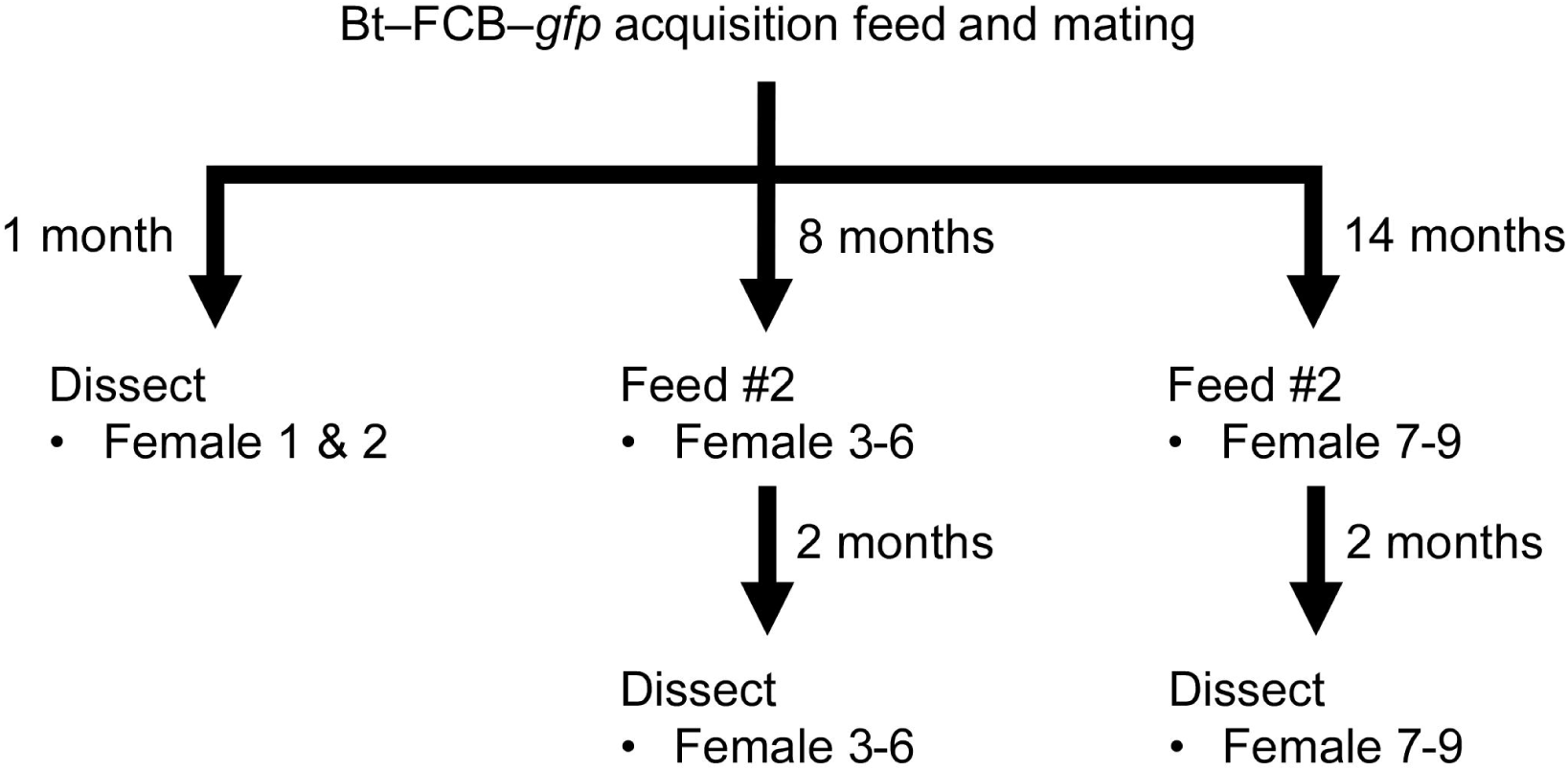
The experimental timeline to infect female *O. turicata–*KS with *Bt–*FCB–*gfp*.

With the remaining ticks, we fed them individually on naïve mice and confirmed infection of *Bt–*FCB–*gfp* in four of seven animals by fluorescent microscopy of the blood (Table 5). The blood meal transitioned females into their second gonotrophic cycle and we assessed tick colonization 10 and 16 months after the initial *Bt–*FCB–*gfp* acquisition (Table 5). By the completion of the second gonotrophic cycle, *Bt–*FCB–*gfp* was visualized in midguts (Table 5), salivary glands (Fig. 8 A–D), and reproductive tissues (Fig. 9 A–D). Colonization in the ovaries was primarily associated with connective tissue, while we could not conclusively detect fluorescing spirochetes in oocytes. We also fed 120 second instar stage nymphal progeny from two infected females (Females 4 and 5 in Table 5) on mice and we failed to detect spirochetes in the mouse blood by microscopy nor did the animals seroconvert to *B. turicatae* protein lysates. These findings indicated that despite successful invasion of the reproductive system by *Bt–* FCB after the second gonotrophic cycle, persistent colonization of oocytes was undetectable.

**Figure 8.**
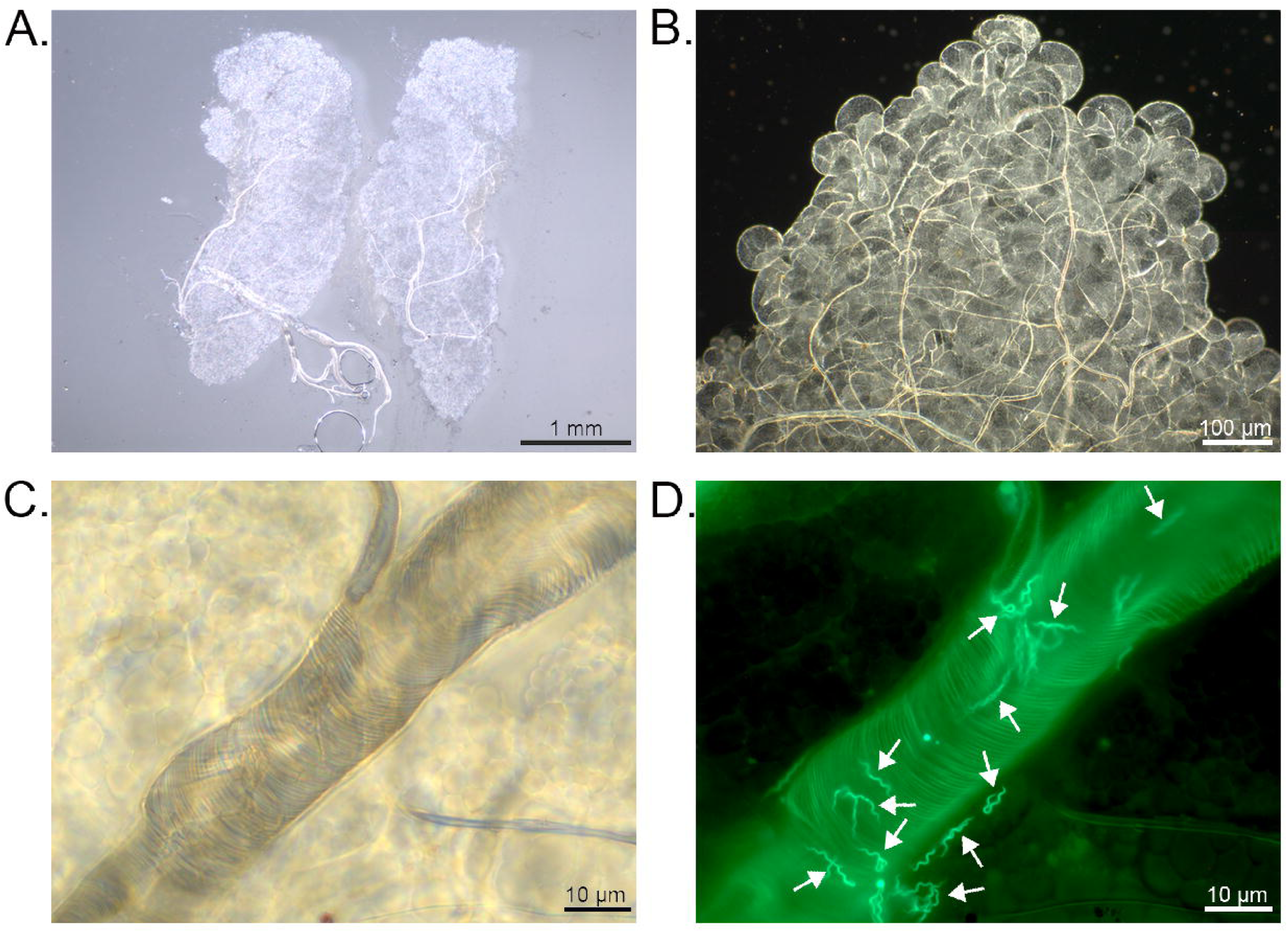
Colonization of salivary glands by FCB-*gfp* spirochetes. (A) shows intact salivary glands excised from an infected *O. turicata* female were gently squashed under a coverslip. (B) shows a single salivary gland lobe (10× objective), with numerous secretory alveoli and confluent alveolar ducts. (C) is a dark-field image (100× oil immersion) centered on an alveolar duct surrounded by glandular tissue. (D) The same field viewed under a GFP filter, revealing abundant fluorescent *Bt–*FCB–*gfp* within the salivary duct and adjacent tissue. Arrowheads point to individual spirochetes. Panels A– D show an example from a single infected female, which represents the remaining six females with microscopically detectable salivary gland infection.

**Figure 9.**
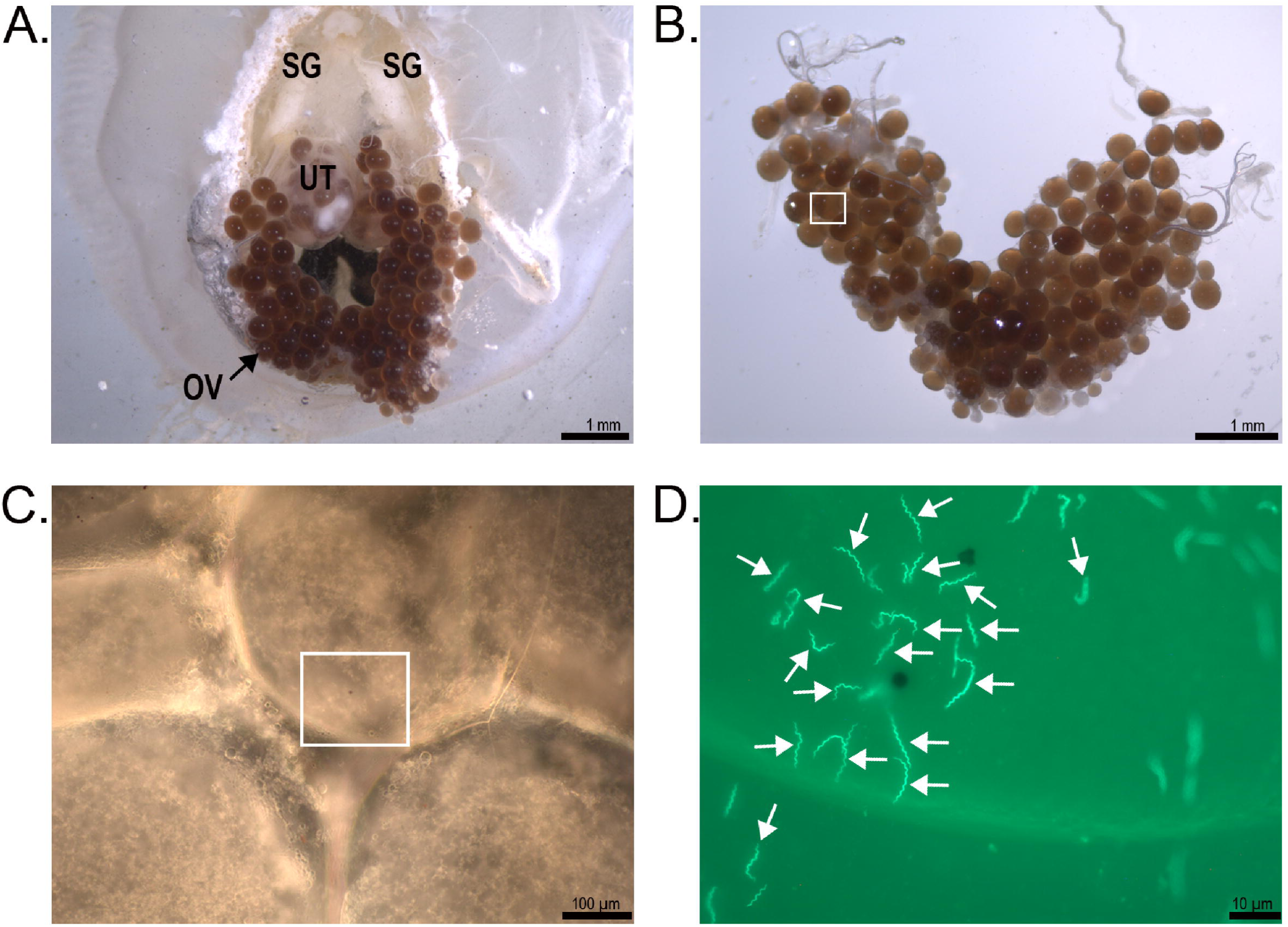
Colonization of tick ovaries by FCB-*gfp* spirochetes. In (A) the dorsal cuticle and midgut tissues were removed from an infected *O. turicata* female, exposing the salivary glands (SG), ovaries (OV), and uterus (UT). In (B), the ovaries were excised and spread on a microscopy slide. In (C), we show a dark-field image (10× objective) showing the area boxed in panel B. In (D) is a 63x oil immersion image of the area boxed in panel C showing fluorescent spirochetes (arrowheads). Panels A–D show an example from a single infected female, which is representative of the remaining four females with microscopically detectable ovary infection.

## Discussion

This study advances our understanding of *B. turicatae* maintenance in *O. turicata*. Male ticks hyperparasitized infected females and acquired *Bt*-SSK1, but not *Bt*-FCB. After digestion and refeeding, both strains remained infectious to mice. However, transovarial transmission (TOT) occurred only with *Bt*-SSK1, which remained infectious through offspring and across a second oviposition within the same gonotrophic cycle. In contrast, progeny from *Bt*-FCB-colonized females were free of spirochetes. Studies using *Bt–*FCB–*gfp* showed that although this strain invaded reproductive tissues, it failed to colonize developing oocytes. These findings underscore the importance of strain–specific differences tick colonization and transmission of *B. turicatae*.

The identification and characterization of hyperparasitism yielded unexpected insights into the biology of *O. turicata* and the transmission dynamics of *B. turicatae*. Although hyperparasitism was once considered rare in *O. turicata* (16), our observations indicated otherwise. Our findings were more consistent with reports for *Ornithodoros parkeri*, *O. hermsi*, and *Ornithodoros erraticus*, in which males commonly feed on engorged conspecifics (9, 10). Interestingly, the differences in the maintenance of *Bt–* SSK1 and *Bt–*FCB in male ticks suggested that the infectious dose may drive colonization after hyperparasitism. Determination of the infectious dose imbibed by male ticks was not feasible because it was discovered post factum and we did not determine spirochete densities imbibed by female ticks. Moreover, male ticks died before we could determine if they were free of *Bt–*FCB or colonized but unable to transmit the strain to naïve mice. While hyperparasitism may influence pathogen prevalence in soft ticks (9, 10), whether it occurs in nature and by other developmental stages of *O. turicata* remains to be investigated.

We used indirect and direct approaches to evaluated TOT in the offspring of infected female *O. turicata*. TOT was indirectly assessed by feeding F1 offspring on naïve mice and assessing infection frequencies in the animals, while qPCR was used to directly determine filial infection rates. Prior to testing F1 offspring, the larva and first two nymphal instar stages were reared on mouse pups. There were several advantages to this approach, including limiting potential human exposure to *B. turicatae*–infected larvae. This was important because Edward Francis reported that a bite from a single larval *O. turicata* was sufficient to infect him (19). Another advantage to testing third instar stage nymphs was the increased likelihood of detecting *B. turicatae* by qPCR in individual ticks. Previously, we have had limited success detecting *B. turicatae* in larva and early instar stage nymphs. However, in the third instar stage and beyond, *B. turicatae* detection is feasible. These findings suggest that *B. turicatae* replicate throughout *O. turicata* development.

A potential limitation of indirectly and directing assessing TOT in third instar stage nymphs was co-feeding transmission of *B. turicatae*. Co-feeding transmission is the ability of uninfected ticks to acquire a pathogen from infected ticks that are engorging on the same naïve animal. We do not think this route of transmission occurred given the rapid feeding behavior of argasid ticks and filial infection frequencies that were detected. For example, several cohorts of third-instar stage nymphs had filial infection frequencies of 5% (1 of 20 ticks). We reasoned that if co-feeding transmission occurred during the larval and first two instar stages, filial infection rates would have been higher. An alternative was to rear ticks individually on mice to determine filial infection frequencies, but this was impractical.

In all cohorts of ticks evaluated, the average frequency of TOT in the first gonotrophic cycle was significantly less compared to the second cycle. Similar trends in TOT were observed for *Borrelia* species vertically transmitted by *O. erraticus* and *Ornithodoros savignyi* (20, 21). Initially, we reasoned that the blood meal in the second gonotrophic cycle may support TBRF spirochete replication and oocycte colonization, but several lines of evidence signaled otherwise.

Our work indicated that dissemination kinetics of *B. turicatae* was a significant driver of oocycte colonization. When we unexpectedly observed gonotrophic dissociation, *Bt–*SSK1 was detected by qPCR in the offspring of two *O. turicata–*KS females after the second oviposition of the first gonotrophic cycle. Importantly, these two females failed to vertically transmit *Bt–*SSK1 in first oviposition. Together, these findings suggest that TOT of *B. turicatae* is a time*–*dependent process in which spirochete dissemination to the reproductive tissues is required before oocyte colonization and subsequent vertical transmission can occur.

Reproductive tissues of the tick impose a strong bottleneck for TOT of microbial organisms and it is important to understand the stage of development and oogenesis when ticks are colonized (22, 23). There are five stages of oogenesis in adult female ticks, and stages I and II occur prior to blood feeding of ixodid and argasid ticks. There are species of argasids, including *O. turicata* (14), in which mating alone activates the progression of oogenesis from stages I and II into the subsequent stages (14, 24). Studies by Abdel-Hamid and colleagues reported that vertical transmission of *Borrelia crocidurae* in *Ornithodoros erraticus* occurs in the first two stages of oogenesis (25). In the study, they used virgin infected females, meaning that the ticks were likely colonized with *B. crocidurae* as nymphs. In our work, the female ticks used were either mated immediately after the *B. turicatae* acquisition bloodmeal or prior to blood feeding. Consequently, oogenesis was activated and all stages were present. Ongoing work is focused on infecting cohorts of *O. turicata* at the nymphal stage and defining TOT throughout oogenesis.

To further understand TOT, we generated *Bt–*FCB that stably expressed *gfp.* This was important because the only other strain of *B. turicatae* that has been transformed is 91E135 (26–30). To generate a *gfp*-expressing 91E135 strain, P*flaB*-*gfp* was targeted for insertion at the 5′ end of the 158 kb linear megaplasmid (29). However, after transformation the cassette recombined to a 40 kb linear plasmid. While spirochetes expressed *gfp* in *O. turicata* for over 18 months (29), the recombination event suggested that the integration site on the megaplasmid was unstable. In our current study transforming *Bt–*FCB, we chose to integrate the P*flaB–gfp* cassette in the intergenic region between *bta019* and *bta020* of the megaplasmid. After transformation, P*flaB–gfp* remained stably integrated. Moreover, when individual *Bt–*FCB–*gfp* female ticks were fed on naïve mice, 57% became infected, which were similar infection frequencies as 91E135–*gfp* (29). This work establishes *Bt*–FCB–*gfp* as a genetically stable and biologically relevant tool to investigate *B. turicatae* persistence and transmission in *O. turicata*.

The inability to detect *Bt*–FCB directly or indirectly in the progeny of spirochete-colonized female ticks prompted us to further examine their reproductive tissues after infection with *Bt*–FCB–*gfp.* We initially thought that *Bt–*FCB was incapable of penetrating and colonizing female reproductive tissues. However, by the completion of the second gonotrophic cycle we detected numerous fluorescing spirochetes in the ovaries of female *O. turicata*. While *Bt–*FCB can invade reproductive tissues it remains unknown if they can penetrate developing oocytes but undergo population reduction. Population reduction has been observed in *Coxiella*-like endosymbionts in *Rhipicephalus* ticks and *Rickettsia* species in whiteflies (31, 32). These bacteria are abundant in early-stages of oogenesis but undergo sharp reductions as oocytes mature. The absence of *Bt*–FCB in progeny ticks may result from a decline in spirochete abundance during oogenesis that prevents persistent colonization of mature oocytes.

Our findings further support prior work that TOT efficiency of TBRF spirochetes varies by strain (33, 34). In 1935 Charles Wheeler evaluated TOT of TBRF spirochetes in 672 progenies from infected *O. hermsi* females and reported a 0.29% filial infection rates (35). At the time of Wheeler’s work, it was unknown that *O. hermsi* could transmit two closely related species of TBRF spirochete (*B. hermsii* and *B. nietonii*), so it is unclear what species was used in his studies. More recently, Schwan and colleagues reported TOT of *B. nietonii* MTW–4 in 84.4% of *O. hermsi* larval cohorts screened (33), indicating strain or species variation. For *B. turicatae,* there have been limited studies in defining vertical transmission of this species because relatively few bacterial strains have been available for experimentation (13).

As we have expanded the number of laboratory *B. turicatae* strains (36), we can now perform functional genomics analyses to identify the molecular mechanisms of TOT. Experimental screening of additional isolates with TOT and non-TOT phenotypes is ongoing. Future pangenome analyses are expected to identify the genomic regions and genetic candidates associated with TOT phenotypes (37, 38). With advances in *B. turicatae* and *O. turicata* genomics (39), the foundation has been established to understand how TBRF spirochetes persist in their tick vectors.

## Materials and Methods

### Ethics Statement

All mice used in this study were Institute of Cancer Research (ICR) strain from a colony maintained at Baylor College of Medicine (BCM). Animal use was reviewed and approved by the BCM Institutional Animal Care and Use Committee (protocol AN*–*7086) and complied with the United States Public Health Service policy and the Guide for the Care and Use of Laboratory Animals.

### Strains of B. turicatae

Non-clonal *B. turicatae* isolates were cultured in Barbour-Stoenner*–*Kelly*–*IIB (BSK-IIB) medium without L*–*cysteine and dithiothreitol (40). The *Bt–*SSK1 strain was isolated from the blood of a mouse after feeding a cohort of transovarially infected second*–*instar stage *O. turicata* nymphs on the animals (14). The *Bt–*FCB strain was isolated from the blood of a dog that originated in Florida (41). *Bt–*SSK1 was passaged twice from the original isolation, while *Bt-*FCB was passaged 12 times from original isolation.

### Tick colonies

Uninfected colonies of *O. turicata* originated from Kansas and Texas and were maintained at Baylor College of Medicine (14, 15, 29, 42). We previously reported that the *O. turicata–*KS colony was free of *B. turicatae* (29, 42), and these ticks were used as a laboratory breeding colony. For *O. turicata–*TX, these females were the F1 progeny of an uninfected field-collected tick that originated from Austin, Texas (14). The progeny were determined to be uninfected with *B. turicatae* by feeding them on mice, collecting blood for 10 consecutive days, and inspecting the blood by dark field microscopy to detect spirochetes. Thirty days after feeding ticks on mice, the animals were exsanguinated and serum was collected for immunoblotting (described below).

### Infecting and breeding *O. turicata* females

To generate experimental groups of infected ticks, five uninfected *O. turicata* females per *B. turicatae* strain underwent an acquisition feed on spirochetemic mice. Mice were infected with *Bt-*SSK1 or *Bt-*FCB by intraperitoneal inoculation with 1 x 10^6^ to 1 x 10^7^ spirochetes in 200 µL of BSK-IIB medium or by tick bite. A drop of blood was collected daily from the tail tip for microscopic confirmation of spirochetemia, and a 2.5 µL aliquot was mixed with 47.5 µL of the SideStep Lysis & Stabiliztion Buffer (Agilent, Santa Clara, CA, USA) and stored at -80°C until analyzed by qPCR to quantify the spirochetemia (see below). Ticks were placed on the shaved abdomen of isoflurane-sedated mice and allowed to feed to repletion.

Freshly fed females were paired with unfed, uninfected *O. turicata* males or kept solitary (ticks that had mated prior to blood feeding) inside ventilated 50- or 15 mL conical tubes (VWR, Radnor, Pennsylvania, USA). Ticks were housed in glass desiccators under standard insectary conditions (24 ± 2 °C, 85-90% relative humidity) (43). Tubes were inspected every two to three weeks. In all experiments eggs were kept separated according to the originating female tick, and resulting larvae were fed on 2-to 5-day-old mouse pups, within two months of hatching. Progeny of experimental females were fed again on mouse pups to reach the second instar stage. Only naïve (unexposed to *B. turicatae*) animals were used for rearing ticks and the pups were never recycled.

### Determination of murine infection after tick feedings

Adult ticks were fed on a naïve mouse individually to assess their *B. turicatae* infection status. To determine whether TOT of spirochetes occurred, a group of 10 to 15 second instar stage nymphs were placed on the shaved abdomen of a sedated mouse and allowed to feed to repletion. Four groups of second instar stage nymphs per cohort of progeny from an individual female (or all progeny if ≤ 60 ticks were available) were tested across two consecutive gonotrophic cycles. The exposed mice were bled by a tail nick on days four, five, nine, and 10 after tick feeding, and slides were prepared using 3–5 µL of fresh, undiluted blood. In each slide, 30 fields of view were examined for spirochetes under a dark field microscope using a 20x objective (Zeiss Axio Imager A2, Zeiss, Munich, Germany). All animals that tested negative by dark field microscopy were assessed for infection by immunoblotting to determine seroconversion.

### Immunoblotting

Four weeks after murine exposure to ticks, animals were terminally bled by cardiac puncture. Six hundred to one mL of whole blood was centrifuged at 6,000 g for 15 minutes, and the serum was collected to determine IgG reactivity against *B. turicatae* protein lysates. SDS–PAGE and immunoblotting was performed as previously described (15). Serum sample dilutions were 1:200 and Rec-protein G-horseradish peroxidase (Life Technologies, Carlsbad, CA, USA) was diluted 1:4,000. Antibody binding to *B. turicatae* protein lysates was determined by chemiluminescence using ECL Western blotting detection reagents (GE Healthcare, Buckinghamshire, UK).

### Spirochetemia quantification in murine blood

Spirochete densities in mouse blood prior to the acquisition blood meal were determined by qPCR using primers and probes for a fragment of the *flaB* gene (Table 6), as previously reported (44). A standard curve of 1 x 10^4^ to 1 x 10^7^ *B. turicatae* spirochetes per mL was generated with *in vitro* grown spirochetes resuspended in PBS-MgCl_2_ and spiked into mouse blood. For each sample, 3 µL of the blood/buffer mixture diluted 10-fold with nuclease-free water was used in a 20 µL reaction, with each reaction run in triplicate. The assays were performed in 384-well plates on a CFX384 Real Time System C1000 Touch Thermal Cycler (Bio-Rad Laboratories, Hercules, CA, USA) using SsoAdvanced Universal Probes Supermix (Bio-Rad).

**Table 6.** Plasmids and strains used in this study.

| Plasmid or Strain | Description<sup>A</sup> | Source |
| --- | --- | --- |
| **Plasmid** |  |  |
| pGEM-T Easy | TA cloning vector; Amp<sup>r</sup> | Promega |
| pUAMS369 | pGEM-T Easy::PflaB-gfp (BamHI/HindIII-flanked); Amp<sup>r</sup> | 18 |
| pJD44 | aph[3']-IIIa marked derivative of pBSV2; Kan<sup>r</sup> | 47 |
| pJD44::gfp | pJD44 containing the PflaB-gfp cassette; Kan<sup>r</sup> | This study |
| pUAMS425 | pGEM-T Easy with 5' and 3' flanking regions for lp158 integration; Amp<sup>r</sup> | This study |
| pBtKan-gfp | pUAMS425 containing the aph[3']-IIIa PflaB-gfp cassette; Kan<sup>r</sup>, Amp<sup>r</sup> | This study |
| **Strain** |  |  |
| *E. coli* |  |  |
| TOP10F' | F' [lacI<sup>q</sup>Tn10(Tet<sup>r</sup>)] mcrA Δ(mrr-hsdRMS-mcrBC) □80lacZΔM15 nupG ΔlacX74 recA1 araΔ139 Δ(ara-leu)7697 galU galK rpsL (Strep<sup>r</sup>) endA1 | Life Technologies |
| *B. turicatae* |  |  |
| Bt-SSK1 | *B. turicatae* strain cultured from blood after feeding a tick on a mouse | 14 |
| Bt-FCB | *B. turicatae* strain FCB, cultured from blood of a dog | 41 |
| Bt-FCB-gfp | BtFCB with gfp-aph[3']-IIIa alleles inserted into the megaplasmid; Kan<sup>r</sup> | This study |

**A:** Amp, ampicillin; Kan, kanamycin

### Molecular detection of *B. turicatae* in ticks

Filial infection rates were determined on a subset of progeny nymphal ticks following their molt to the third instar stage. Genomic DNA (gDNA) was individually extracted from 20 randomly selected whole ticks per cohort using the DNeasy Blood and Tissue kit (Qiagen, Hilden, Germany). Ticks from the same cohort were extracted on the same day using the same batch of reagents. Extraction controls included a blank sample (DNeasy reagents only), two uninfected and one *B. turicatae*-infected *O. turicata* ticks to account for potential carry-over contamination and reaction inhibition. To avoid cross-contamination, each cohort was assessed for *B. turicatae* infection in a dedicated 96-well plate using a duplex qPCR assay, as previously described (15). Primer and probes were designed to *B. turicatae flaB* and the *O. turicata* β*-actin* genes (Table 6). A total of 25 ng of tick gDNA was added to a 20 µL reaction containing Brilliant II qPCR Mastermix (Agilent, Santa Clara, CA, USA), 400 nm of primers, 300nM of probes, 30 nM reference dye, and nuclease-free water. Each sample was run in triplicate. Each assay included standard curves prepared with *flaB* and *B-actin* plasmids (1 x 10^1^ to 1 x 10^6^ plasmid copies per 2.5 µL) (15), tick extraction controls, and nuclease-free water as a control. The assays were performed on a QuantStudio™ 7 Pro Real-Time PCR System (Applied Biosystems, Foster City, CA, USA).

### *Bt–*FCB genetics

Bacterial strains and plasmids used in this study are listed in Table 6. *Escherichia coli* strain Top10F’ (Life Technologies, Carlsbad, CA) was used for cloning and plasmid propagation. *E. coli* was grown in lysogeny broth (LB) supplemented with 100 µg/ml ampicillin or 50 µg/ml kanamycin when necessary. *Bt-*FCB was passaged no more than twice beyond the original frozen stock and cultured at 35°C with 3% CO_2_ in modified BSK (mBSK) medium with 12% rabbit serum at pH 7.6 unless noted otherwise (45, 46). mBSK was supplemented with 150 μg/ml kanamycin when appropriate. Primers used in this study are listed in Table 7. Plasmid and genomic DNA (gDNA) were isolated using the Wizard Plus SV Miniprep DNA Purification System (Promega Corp., Fitchburg, WI), and PCR amplifications for cloning were performed with high-fidelity PrimeSTAR Max DNA Polymerase (TaKaRa Bio, Mountain View, CA). All plasmids and cloned fragments were Sanger sequenced to confirm that no mutations were introduced during cloning.

**TABLE 7.**
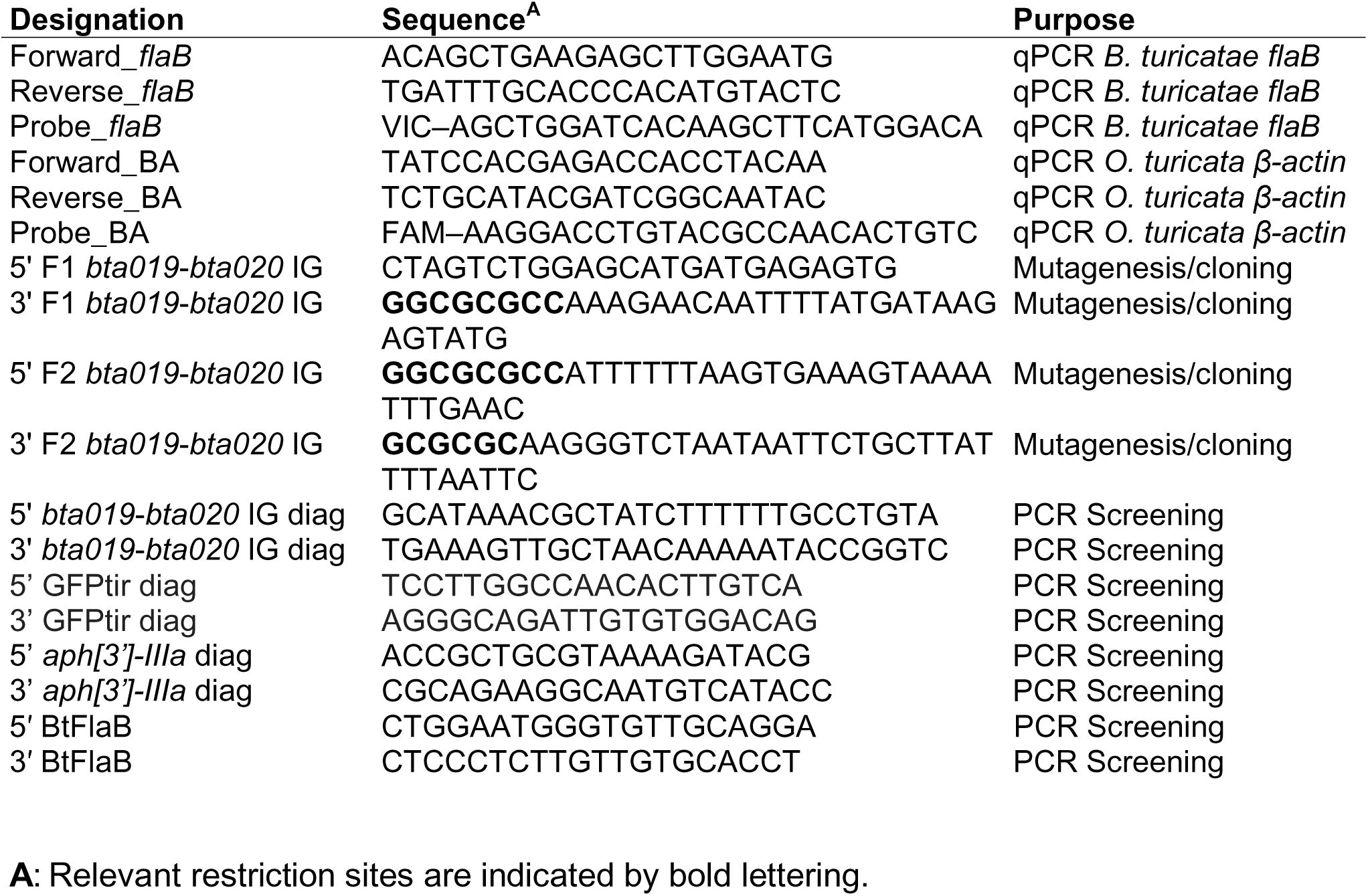
Primers and probe used in this study.

To generate a *gfp-*expressing isolate of strain FCB, the P*flaB*-*gfp* reporter developed for relapsing fever *Borrelia* was then excised from pUAMS369 with *Bam*HI and *Hind*III and ligated into the MCS of the *B. burgdorferi* shuttle vector pJD44, generating the shuttle vector pJD44::*gfp* (18, 47). Next, a site-specific integration construct targeting the intergenic region between *bta019* and *bta020* on the 158-kb linear megaplasmid was created. 5’ (primers: 5’ F1 *bta019*-*bta020* IG and 3’ F1 *bta019*-*bta020* IG) and 3’ (primers: 5’ F2 *bta019*-*bta020* IG and 3’ F2 *bta019*-*bta020* IG) regions flanking the integration site (1016- and 1003-bp in size, respectively) were amplified and TA-cloned into pGEM-T Easy. The flanking regions were then ligated together, with an AscI restriction site between them, into pGEM-T Easy, generating the plasmid pUAMS425. The *aph*[3′]-*IIIa* kanamycin resistance marker and P*flaB*-*gfp* cassette were excised from pJD44::*gfp* with *Asc*I and ligated into *Asc*I-digested pUAMS425, to generate pBtKan-*gfp*.

Competent cell preparation and electroporation of *Bt*-FCB were performed as previously described (30, 45). Transformants were selected with kanamycin, confirmed by PCR, and the products were separated by electrophoresis in a 0.8% agarose gel and visualized with ethidium bromide staining. Integration of *aph*[3′]-*IIIa-gfp* into the 158-kb linear megaplasmid of BtFCB-*gfp* was confirmed by PCR to amplify internal regions of *gfp* (primers: 5’ GFPtir diag and 3’ GFPtir diag), *aph*[3′]-*IIIa* (primers: 5’ *aph*[3’]-*IIIa* diag and 3’ *aph*[3’]-*IIIa* diag), and *flaB* (primers: 5’ BtFlaB and 3’ BtFlaB). PCR was also performed to amplify across the insertion site between *bta019* and *bta020* (primers: 5’ *bta019*-*bta020* IG diag and 3’ *bta019*-*bta020* IG diag). GeneRuler DNA Ladder Mix (ThermoFisher Scientific, Waltham, MA) served as the molecular weight standard.

Imaging of *gfp-*expressing bacteria was performed as previously described (27). Briefly, strains of interest were grown to mid-exponential phase, washed twice with phosphate-buffered saline (PBS)-MgCl_2_, spotted onto a 1% agarose pad, and covered with a coverslip (48). Brightfield and fluorescent images were then captured using an Olympus BX51 microscope with a 100x oil-immersion objective (Olympus, Melville, NY, USA). The bacteria were stored frozen in mBSK with 20% glycerol at -80°C, and cultures were initiated by inoculating four milliliters of mBSK with a scoop of the frozen stock. All cultures were incubated at 35°C in 5% CO_2_ atmosphere and grown to the mid-log phase.

### Assessment of *Bt–*FCB–*gfp* in ticks

A cohort of *O. turicata–*KS females (n=10) were infected with *Bt–*FCB–*gfp* and evaluated at time points shown in Figure 7. To infect ticks, two mice were infected by intraperitoneal injection of ∼ 1 x 10^7^ *Bt–*FCB–*gfp* spirochetes. To avoid hyperparasitism, freshly fed females were paired with males that fed on the same animals. Two females were dissected at one-month post-acquisition (Fig. 7). The remaining seven females were individually fed on mice to assess the transmission to mice as described above and dissected after completing their second gonotrophic cycle (Fig. 7). Ticks were glued dorsal side up to a bottom of a 60 x 15 mm plastic Petri dishes (VWR, Radnor, Pennsylvania, USA) and chilled for a few hours at 4 °C. The plates were flooded with cold Dulbecco′s Phosphate Buffered Saline (DPBS; Sigma-Aldrich, St Louis, MO, USA) and ticks were dissected under an Axio Stemi stereomicroscope (Zeiss Munich, Germany). Individual tissues were excised, rinsed in DPBS, placed on a slide in a drop of buffer, gently squashed with a coverslip, and examined for spirochetes using an Axio Imager A2 (Zeiss) fluorescence microscope. Images were captured and processed using the ZEN 2012 digital imaging software (Zeiss). Additionally, two cohorts of progeny resulting from the second gonotrophic cycle were tested for TOT by feeding them on mice at the second instar stage, as described above.

### Statistical analyses

To ensure that the experimental system remained statistically robust even when filial infection rates were low (≤5%), we used the binomial formula to calculate the probability of exposure to spirochetes for mice bitten by varying numbers of ticks (49). At 10–15 ticks per animal, the probability of at least one infected tick successfully feeding remained high, even under low-filial infection rates. All statistical analyses were performed in R version 4.4.1 (R Core Team 2020). We calculated 95% confidence intervals for the point estimates of filial infection rates obtained with qPCR for each progeny cohort using the Wilson interval with continuity correction, as implemented in the *BinomCI()* function from the *DescTools* package (50). Filial infection rates between the two gonotrophic cycles were compared using the Fisher’s exact test (α=0.05) and the Cliff’s Delta (δ) from the *effsize* package (51), to test for statistical significance, as well as the magnitude and direction of the observed differences. The later metric is suitable for analysis of non-normally distributed binary data and ranges from -1 to 1. A δ of 0 indicates no difference between groups, positive values indicate higher measurements in the first group compared to the second, and negative values indicate lower measurements in the first group compared to the second. Effect size is interpreted as follows: (δ) < 0.15: Negligible; 0.15–<0.33: Small; 0.33–<0.47: Medium; ≥ 0.47: Large. (52). Fisher’s exact test and Cliff’s delta were calculated based on the total number of screened ticks across gonotrophic cycle 1 and 2; per-female Fisher’s tests were also performed and were consistent with the pooled results.

## Acknowledgments

We thank Brittany Armstrong for providing a critical review of this manuscript. This work was supported by grants AI191624 and AI197410 from the National Institute of Allergy and Infectious Diseases, National Institutes of Health, The Texas EcoLabs, and the Lyda Hill Philanthropies. The funders had no role in study design, data collection and interpretation, or the decision to submit the work for publication. We thank Tom Schwan for originally providing the *B. turicatae* FCB strain.

